# Predicting Risk of Transfusion-Induced Red Blood Cell Alloimmunization Using Statistical and Machine Learning Approaches in the Recipient Epidemiology and Donor Evaluation Study (REDS-III) Database

**DOI:** 10.1101/2025.09.21.677587

**Authors:** Duan D. Wang, Lingbo Zhou, Jiawen Chen, Jeanne E. Hendrickson, Matthew S. Karafin, Rui Xiao

**Affiliations:** North Carolina School of Science and Mathematics - Durham, Durham, NC; Department of Biostatistics, University of North Carolina at Chapel Hill, Chapel Hill, NC; Department of Laboratory Medicine, Yale University School of Medicine, New Haven, CT; Department of Pathology and Laboratory Medicine, Emory University School of Medicine, Atlanta, GA; Department of Pathology and Laboratory Medicine, University of North Carolina at Chapel Hill, Chapel Hill, NC; Department of Biostatistics, Epidemiology and Informatics, Perelman School of Medicine, University of Pennsylvania, Philadelphia, PA

**Keywords:** alloimmunization, blood transfusion, prediction, XGBoost, large language model

## Abstract

Red blood cell (RBC) alloimmunization is a common complication from blood transfusion, often resulting in accelerated donor RBC destruction. Patients show substantial variation in their predisposition to RBC alloimmunization. Previous studies have identified several risk factors, but to our knowledge, there have been no studies that predict risk of RBC alloimmunization by modeling multiple potential risk factors simultaneously. Here, our study represents the first attempt to build prediction models for RBC alloimmunization using the large sample size and rich set of potential risk factors available in the Recipient Epidemiology and Donor Evaluation Study (REDS-III) recipient database. To develop the prediction models, we applied a range of approaches, including traditional statistical models (logistic regression), and modern machine learning (including gradient boosting, random forest, and XGBoost), deep learning (the multilayer perceptron method), and large language models (LLM). XGBoost demonstrates the overall best performance among models providing uncertainty quantification (F1= 0.672 and area under the ROC curve [AUC-ROC]=0.752). LLMs show promising results with the best F1 scores (0.677-0.687), though they are limited by their inability to provide uncertainty estimates, they hold the potential for use as an interactive chatbot for patients. Although there is ample room for performance improvement, limiting the analysis to patients predicted with >80% confidence by XGBoost resulted in a substantially improved AUC-ROC of 0.919, which can be of potential clinical significance.

## Introduction

Red blood cell (RBC) alloimmunization, also known as the formation of antibodies against non-self antigens on RBCs, can occur after exposure to RBC through blood transfusion or pregnancy. While most of the common immediate adverse effects of RBC transfusion, including fever, chills and urticaria, are not highly consequential, certain acute and delayed reactions such as hemolytic transfusion reactions (HTR) can be fatal (Arthur and Stowell 2023).

The prevalence of RBC alloimmunization is 2-5% in the general population (Linder and Chou 2021), but is higher among various groups, including pregnant women (estimated to be as high as 2-12% in Sub-Saharan Africans (Mbalibulha et al. 2015; Natukunda et al. 2011; Osaro and Charles 2010), and 1.15-1.5% in the United States (Geifman-Holtzman et al. 1997; Sugrue et al. 2024)), individuals with sickle cell disease (with reported high estimates of 14.4% (Miller et al. 2013), 16.1% (Mijovic, Perera, and Thein 2013) and even up to 76% (da Cunha Gomes et al. 2019)), and more generally among polytransfused patients, particularly with existing blood disorders such as thalassemias (with estimated prevalence ranging from 7.5% (Cruz Rde et al. 2011) to 33.9% (Thedsawad, Taka, and Wanachiwanawin 2019)).

There are many distinct RBC antigens that can lead to RBC alloimmunization. For instance, the Rhesus (Rh) blood group system, a highly polymorphic and immunogenic system, encompasses multiple Rhesus erythrocyte antigens including C or c antigen, E or e antigen, and D antigen (Avent and Reid 2000).

Since RBC transfusion is among the most common medical interventions in hospitalized patients, better understanding of transfusion-induced RBC alloimmunization is of important clinical significance, for example, allowing targeted preventative strategies in higher-risk groups focusing on the influential and clinically actionable risk factors (Tormey and Hendrickson 2019). The mechanisms underlying RBC alloimmunization, however, have emerged as complex and remain largely elusive. Multiple distinct mechanisms governed by different immune pathways have been hypothesized, depending on the alloantigens that trigger the immune response (Arthur and Stowell 2023). Numerous risk factors can contribute to the development of RBC alloantibody after blood transfusion, including factors from both the donor and transfusion recipient. For instance, RBCs from female donors have been found (Kanias et al. 2017) to be less susceptible to storage and stress hemolysis than those from male donors, and donors with underlying diseases such as thalassemia trait, G6PD deficiency, and sickle cell trait have also been reported to negatively affect transfusion efficacy (Francis et al. 2013; Karafin and Francis 2019; Noulsri and Lerdwana 2023).

Multiple recipient risk factors have been established by previous studies. For example, Karafin et al. (Karafin et al. 2018) found RhD negative status, female sex, and pre-existing diseases such as systemic lupus erythematosus (SLE), sickle cell trait, and sickle cell disease were all associated with a significantly increased risk of alloimmunization. In contrast, Asian race and younger recipient age were associated with lower alloimmunization risk. Other studies have reported additional risk factors, including HLA interactions (Chiaroni et al. 2006; Verduin et al. 2016), inflammation (Hendrickson et al. 2006; Fasano et al. 2015; Evers et al. 2016), the presence of autoantibodies (Arthur and Stowell 2023), and specific genetic profiles (Williams et al. 2018; Sun et al. 2024; Meinderts et al. 2019). Despite these studies, few prediction models exist, particularly those that leverage modern machine learning and foundation models, and those that simultaneously consider multiple potential risk factors are still lacking. In this work, we aim to fill this knowledge gap by building models that predict RBC alloimmunization using a comprehensive list of potential recipient risk factors. To do this, we employ a battery of statistical, machine learning, deep learning, and large language model (LLM) methods including logistic regression, random forest, gradient boosting, XGBoost (Chen and Guestrin 2016), multilayer perceptron (MLP) (Murtagh 1991), and Qwen LLM (Bai et al. 2023).

## Methods

### Overview

As illustrated in **Figure 1**, we first downloaded the REDS-III recipient data and processed the data following the methods described in Karafin et al. 2018 (Karafin et al. 2018). We performed logistic regression analyses to validate association between RBC alloimmunization and the previously reported risk factors in the literature. We then focused on building prediction models accommodating all the potential predictors simultaneously in the training dataset with various statistical, machine learning, deep learning and LLM methods, and assessed performance of the prediction models using 10-fold cross-validation. Cross-validation is a widely used technique to evaluate model performance. In *k*-fold (here *k* = 10) cross-validation, data are randomly split into *k* equal partitions and we build the models k times. Each time one of the k partitions is kept for testing and the other *k* - 1 are used to train the model. Performance is assessed in the testing sample, which includes every data point across the *k* times, by comparing predicted to the observed outcomes.

**Figure 1.**
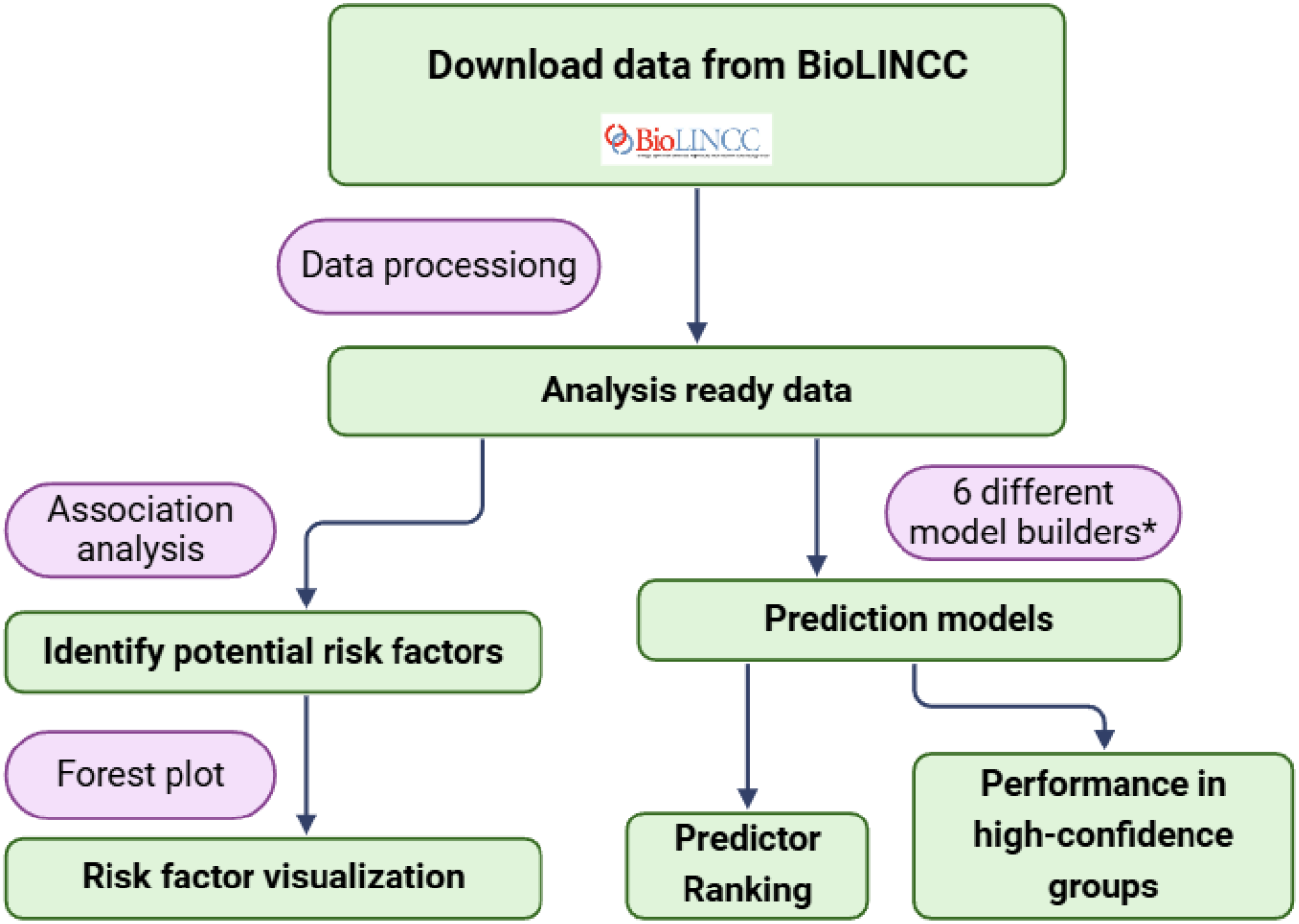
Study overview. * 6 model builders include logistic regression, gradient boosting, random forest, XGBoost, MLP, and Qwen LLM.

### The REDS-III Database

The REDS-III database was developed by the National Heart, Lung, and Blood Institute (NHLBI). REDS-III is a transfusion recipient database that collected electronic health records from four major blood centers, as well as 12 community and academic hospitals in the United States from 2013-2016, including both inpatient and outpatient records. Details of the REDS-III recipient database have been described previously (Karafin et al. 2018; Karafin et al. 2017). We obtained the data through NHLBI’s Biologic Specimen and Data Repository Information Coordinating Center (BioLINCC). The data contains medical records for 637,109 individuals.

### Data Processing

We processed the data from BioLINCC following our previous study (Karafin et al. 2018). In particular, we defined RBC alloantibody responders (for brevity, hereafter referred to as responders) as individuals with at least one clinically significant RBC alloantibody detected at any point after blood transfusion; and defined non-responders as individuals who had a prior RBC transfusion but no clinically significant RBC alloantibody was ever detected during the study period. After data processing, we identified 4,185 responders and 95,847 non-responders for inclusion in the analysis (**Figure 2**). Note that our number of non-responders is larger than previously reported (Karafin et al. 2018), reflecting the differences between the original and public use dataset. For the public use dataset, we modified the criterion of “a negative RBC antibody screen at least 15 days after RBC transfusion” to a more inclusive definition of “a negative RBC antibody screen the same or a later month compared to RBC transfusion”.

**Figure 2.**
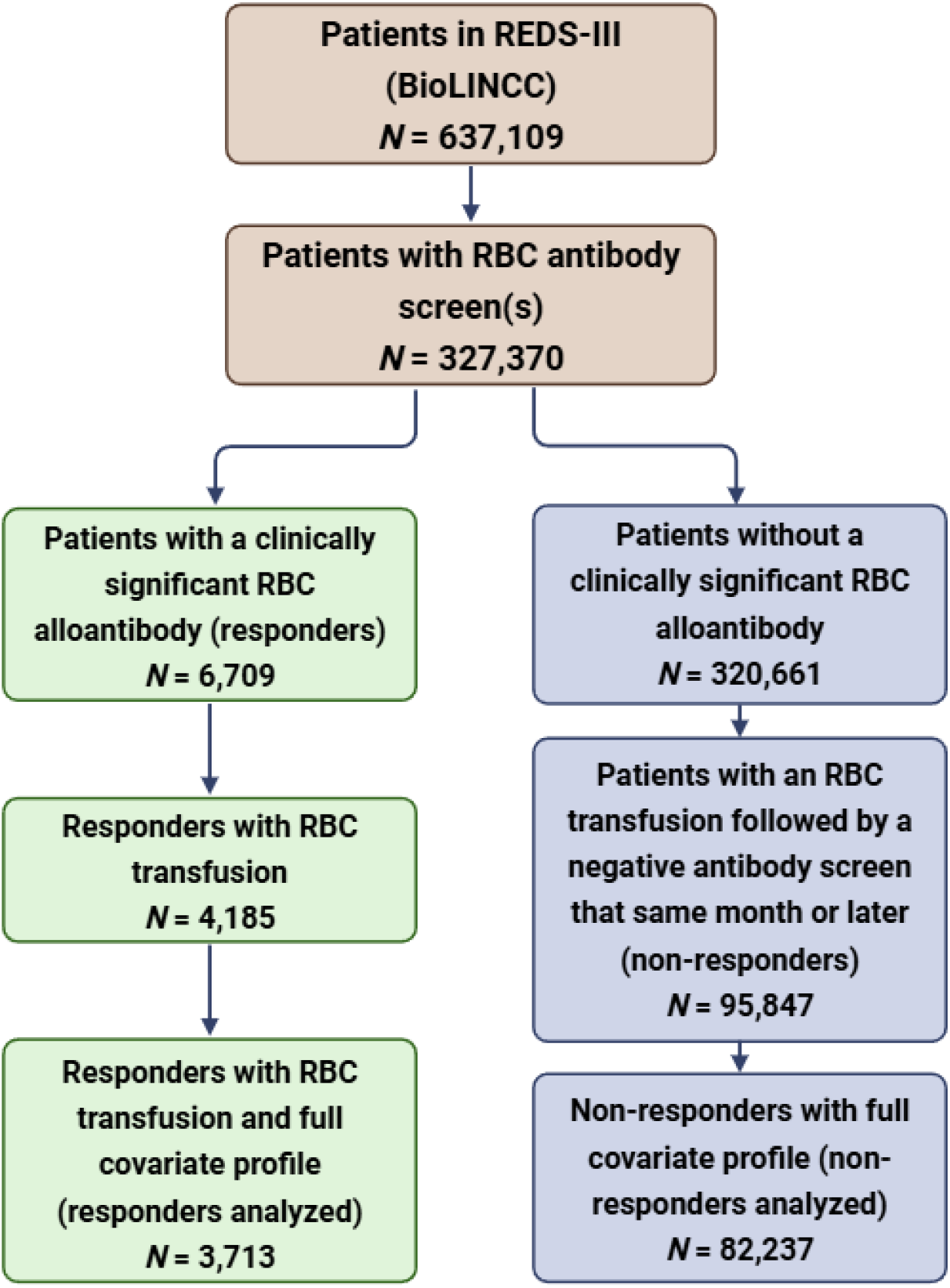
REDS-III sample size and data processing.

We considered the following potential recipient risk factors: age, sex, RhD status, self-reported race, various diseases and pre-existing conditions including leukemia, SLE, malignancy, myelodysplastic syndrome, rheumatoid arthritis, sickle cell disease, sickle cell trait, solid organ transplant, and stem cell transplant. For age, we used similar age categories as in our previous study (Karafin et al. 2018). Disease statuses were based on ICD9 and ICD10 codes (detailed in **Supplementary Table 1**). After excluding individuals with incomplete covariate information, 3,713 responders and 82,237 non-responders remained in our analyses (**Figure 2**).

### Association analysis

We performed logistic regression analysis using *glm* function in R to assess the association between responder status (responded vs. not responded) and each potential risk factor, with results reported as odds ratios (ORs) with 95% confidence intervals (CIs). In the analysis, age was grouped into 10-year categories, following our previous study (Karafin et al. 2018).

### Prediction model development for RBC alloimmunization

We applied four categories of approaches for the development of prediction model, including the classical statistical method logistic regression, three machine learning methods (random forest, gradient boosting, and XGBoost [Chen and Guestrin 2016]), a deep learning method (MLP, Popescu et al. 2009), and a LLM (Qwen, Bai et al. 2023). Logistic regression, random forest and gradient boosting models were trained and evaluated using Python package *sklearn*. XGboost boost models were fit in R using the *xgboost* package. Qwen LLM models with 1.8 billion (Qwen-1.8B) or 7 billion parameters (Qwen-7B) were fine-tuned using Python package *LLaMA-Factory* with both low-rank adaptation (LoRA, Hu et al. 2021) and quantized low-rank adaptation (QloRA, Dettmers et al. 2024). Both LoRA and QloRA allow fine-tuning foundation models (that often use huge numbers of parameters, in the magnitude of billions or even trillions) with a much smaller number of parameters to address the specific research question of interest, in our case, prediction of responders versus non-responders. We obtained Qwen models through Huggingface at: https://huggingface.co/Qwen.

One of the challenges in building the prediction models was the imbalanced responder-to-non-responder ratio (1:22) in this study. To address this, we adopted the Synthetic Minority Oversampling Technique (SMOTE) (Chawla et al. 2002) data augmentation approach, implemented in the Python *imblearn* library, to perform the responder oversampling. The augmented data were then used for all prediction methods except XGBoost, for which we utilized the built-in *scale_pos_weight* option in the *xgboost* R package.

We compared the performance of the prediction models using two metrics: the F1 score and the area under the ROC curve (AUC-ROC). The F1 score combines precision and recall into one metric by calculating a harmonic mean between the two, and therefore, considers the false positive and false negative predictions simultaneously. AUC-ROC, another single-number metric for models that classify a binary outcome (in our context, responder or non-responder), quantifies models’ *overall* capability to distinguish between responders and non-responders across all possible classification thresholds. Both F1 score and AUC-ROC range from 0 to 1 with 1 being the best.

In addition, we ranked the relative importance of the potential predictors for the top performing models. Specifically, for XGBoost models, we used the *importance* function in the *xgboost* R package. For logistic regression and MLP models, we used the *permutation_importance* in the *sklearn*.*inspection* Python package. We then constructed composite importance scores for the top predictors based on the importance ranking from the three methods, weighted by their F1 scores.

## Results

### Demographics of REDS-III participants analyzed

We summarized the basic demographic profiles of the 85,950 REDS-III participants analyzed in this study (**Table 1**). We have an roughly equal numbers of males (49%) and females (51%). Most transfusion recipients (∼88%) were older than 40 years old. Most were self-reported as Caucasian/white (80%).

**Table 1.**
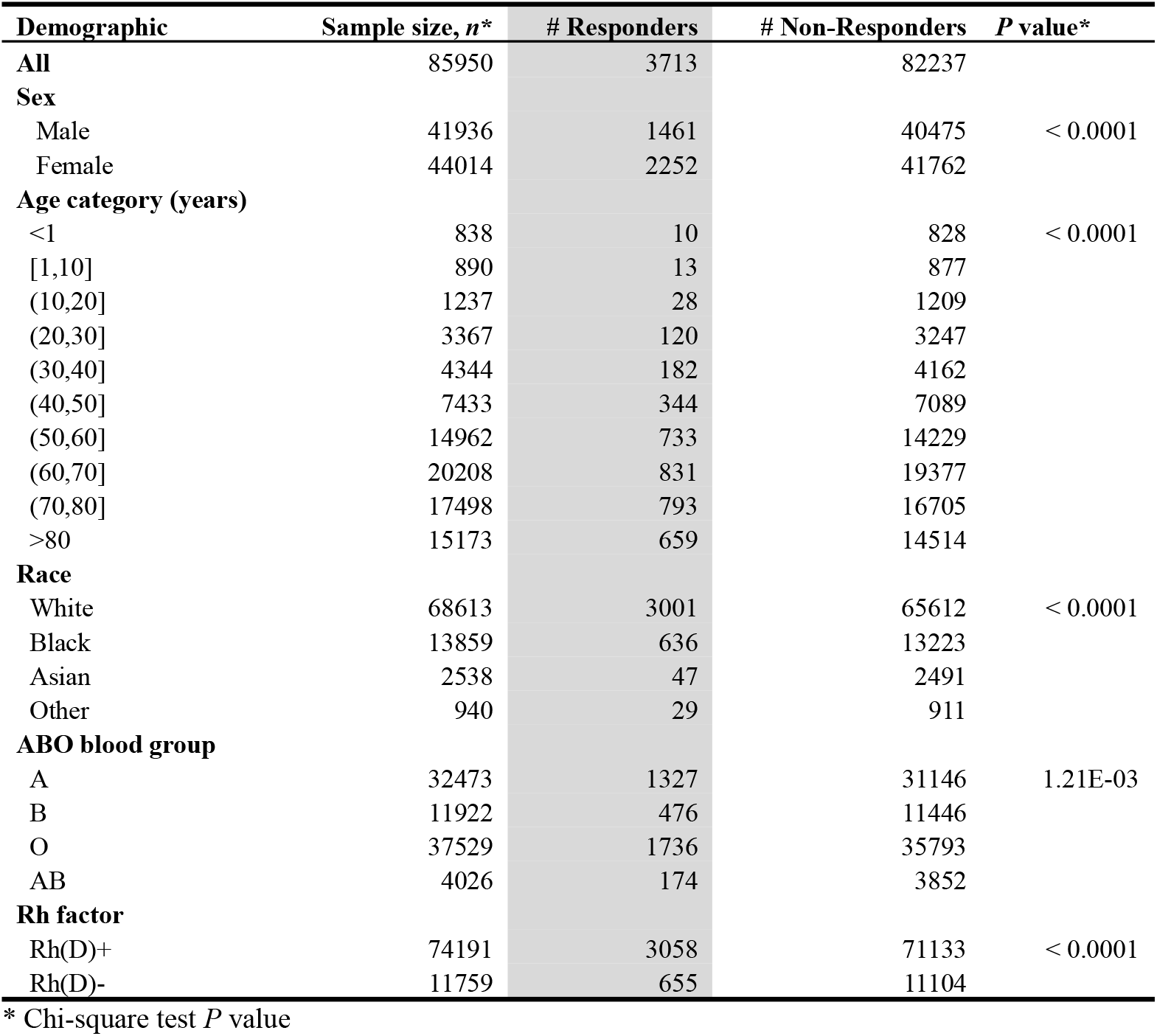
Demographic characteristics of participants analyzed.

| **Demographic** | **Sample size, *n*\*** | **# Responders** | **# Non-Responders** | ***P* value\*** |
| --- | --- | --- | --- | --- |
| **All** | 85950 | 3713 | 82237 |  |
| **Sex** |  |  |  |  |
| Male | 41936 | 1461 | 40475 | < 0.0001 |
| Female | 44014 | 2252 | 41762 |  |
| **Age category (years)** |  |  |  |  |
| <1 | 838 | 10 | 828 | < 0.0001 |
| [1,10] | 890 | 13 | 877 |  |
| (10,20] | 1237 | 28 | 1209 |  |
| (20,30] | 3367 | 120 | 3247 |  |
| (30,40] | 4344 | 182 | 4162 |  |
| (40,50] | 7433 | 344 | 7089 |  |
| (50,60] | 14962 | 733 | 14229 |  |
| (60,70] | 20208 | 831 | 19377 |  |
| (70,80] | 17498 | 793 | 16705 |  |
| >80 | 15173 | 659 | 14514 |  |
| **Race** |  |  |  |  |
| White | 68613 | 3001 | 65612 | < 0.0001 |
| Black | 13859 | 636 | 13223 |  |
| Asian | 2538 | 47 | 2491 |  |
| Other | 940 | 29 | 911 |  |
| **ABO blood group** |  |  |  |  |
| A | 32473 | 1327 | 31146 | 1.21E-03 |
| B | 11922 | 476 | 11446 |  |
| O | 37529 | 1736 | 35793 |  |
| AB | 4026 | 174 | 3852 |  |
| **Rh factor** |  |  |  |  |
| Rh(D)+ | 74191 | 3058 | 71133 | < 0.0001 |
| Rh(D)- | 11759 | 655 | 11104 |  |

\* Chi-square test *P* value

### Association

Consistent with our previous findings (Karafin et al. 2018), logistic regression analyses (summarized in Supplementary section **Association Details** and **Supplementary Figure 1**) confirmed multiple previously reported risk factors, including recipient RhD blood type, ABO blood group, sex, race, age, and multiple diseases such as leukemia, SLE, myelodysplastic syndrome, sickle cell disease and solid organ transplant.

### Prediction performance

According to the F1 score (**Figure 3** top panel), the four Qwen LLMs performed the best, particularly the Qwen-7B models, with F1 score of 0.686 and 0.687 respectively. LoRA or QloRA fine-tuning show similar performance, with a difference of merely 0.001 for both the (more complex with a larger number of parameters) 7B and (less complex with a smaller number of parameters) 1.8B models. Among the other models, logistic regression and XGboost performed similarly, with an F1 score of 0.671 and 0.672 respectively.

**Figure 3.**
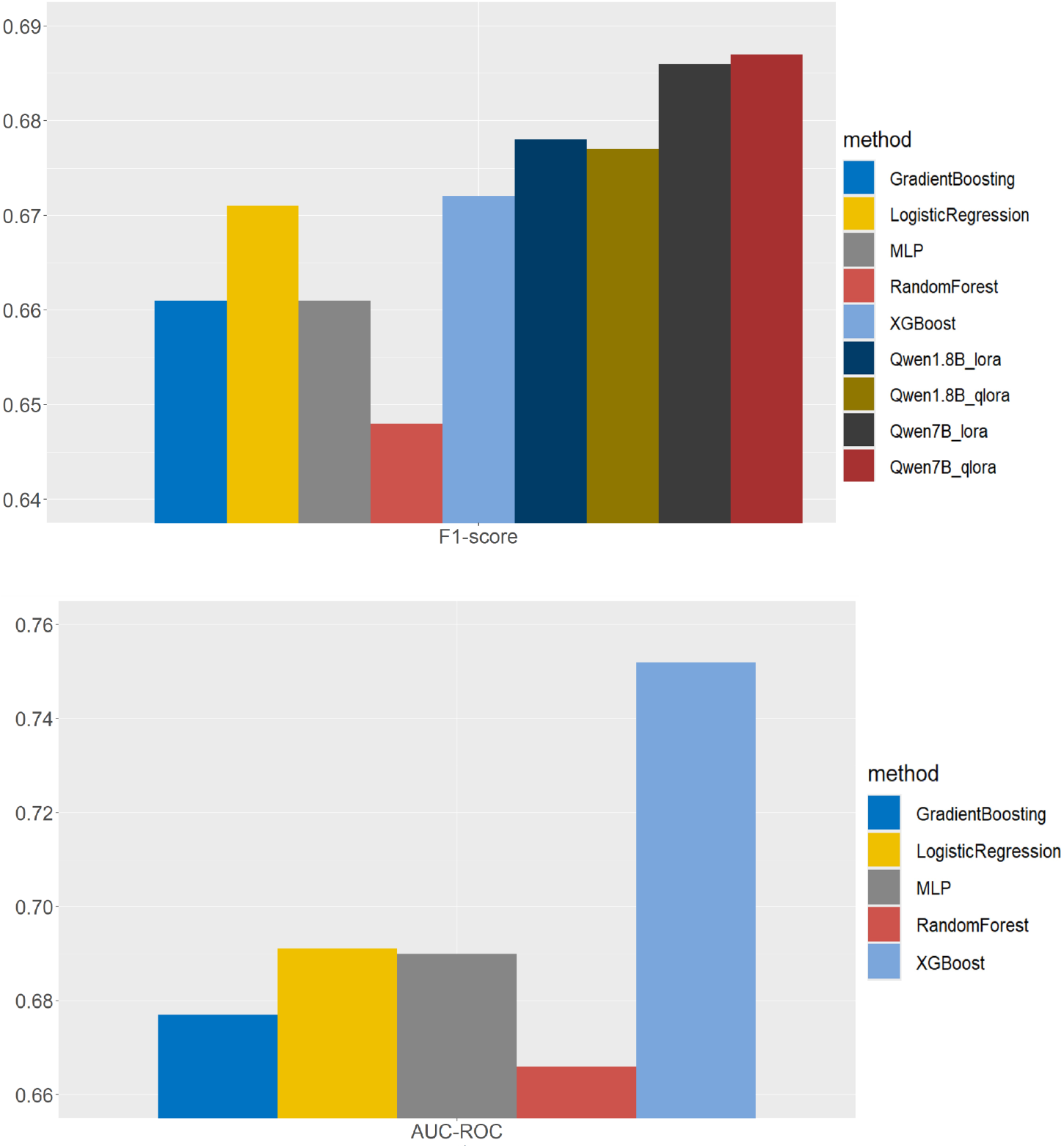
Performance comparison of prediction models. F1-score and area of under the ROC curve are calculated and compared across the methods. Note that Qwen doesn’t provide prediction uncertainty, so does not allow AUC-ROC calculation.

Unlike the other five methods which provide the estimated probability of being a responder, Qwen, as a LLM, only outputs binary (yes or no) predictions. Therefore, we were unable to calculate AUC-ROC for the Qwen models. Among the other methods, XGBoost performed the best, achieving an AUC-ROC of 0.752 compared with 0.666-0.691 for the other four methods (**Figure 3** bottom panel). Given its superior performance among the methods that provide quantitative probabilities for estimating prediction uncertainties, we focus below on the XGBoost model and its variations.

### Prediction performance by sex

As females are known to be at higher risk of developing alloimmunization after transfusion (Karafin et al. 2018), we evaluated the prediction performance separately for male and female participants. We observe similar patterns in the distributions of predicted probability of being a responder (**Supplementary Figure 2**), with actual responders exhibiting the highest peaks near 1 for both sexes. Consistent with the better separation among females, F1 score is higher in females (0.701) compared with males (0.618).

### Prediction performance for patients predicted with high confidence

We note here that the current models are still suboptimal, as the best F1 scores and AUC-ROC are far from the perfect score of 1. Based on **Figure 4** and **Supplementary Figure 2**, most individuals have a predicted probability of being a responder that lies far from 0 or 1. We performed subgroup analysis by focusing only on individuals with predicted probability close to 0 or 1 (e.g., > 0.8 or < 0.2). In this subgroup, the AUC-ROC reached 0.919, highlighting the model’s potential to identify individuals at the highest or lowest risk.

**Figure 4.**
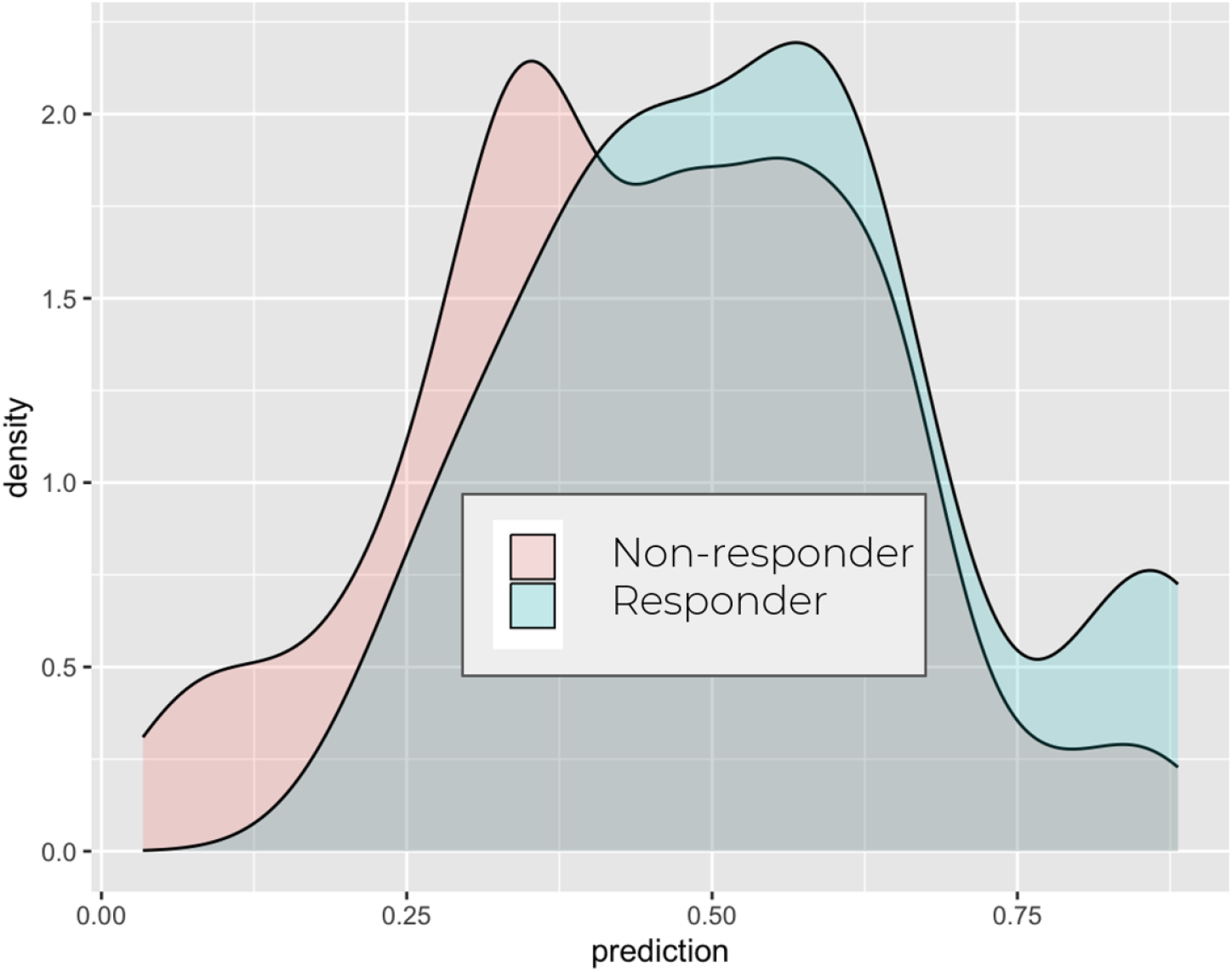
Performance of XGBoost for samples with high confidence predictions. XGBoost model performance for males and females. Density plot for XGBoost predicted probability of being a responder is shown separately for responders (cyan) and non-responders (red).

### Key features driving prediction performance

We also ranked the importance of the predictors. As summarized in **Table 2** across three top-performing models and in **Figure 5** for XGBoost, the top three most influential predictors are age, sex and sickle cell disease, followed by myelodysplastic syndrome, RhD status, Asian race and lupus, in slightly different but largely consistent orders across the three top-performing models, namely XGBoost, logistic regression and MLP. Dropping the top-ranking features substantially impaired the performance of the models as shown in **Supplementary Figures 3 and 4**.

**Table 2.**
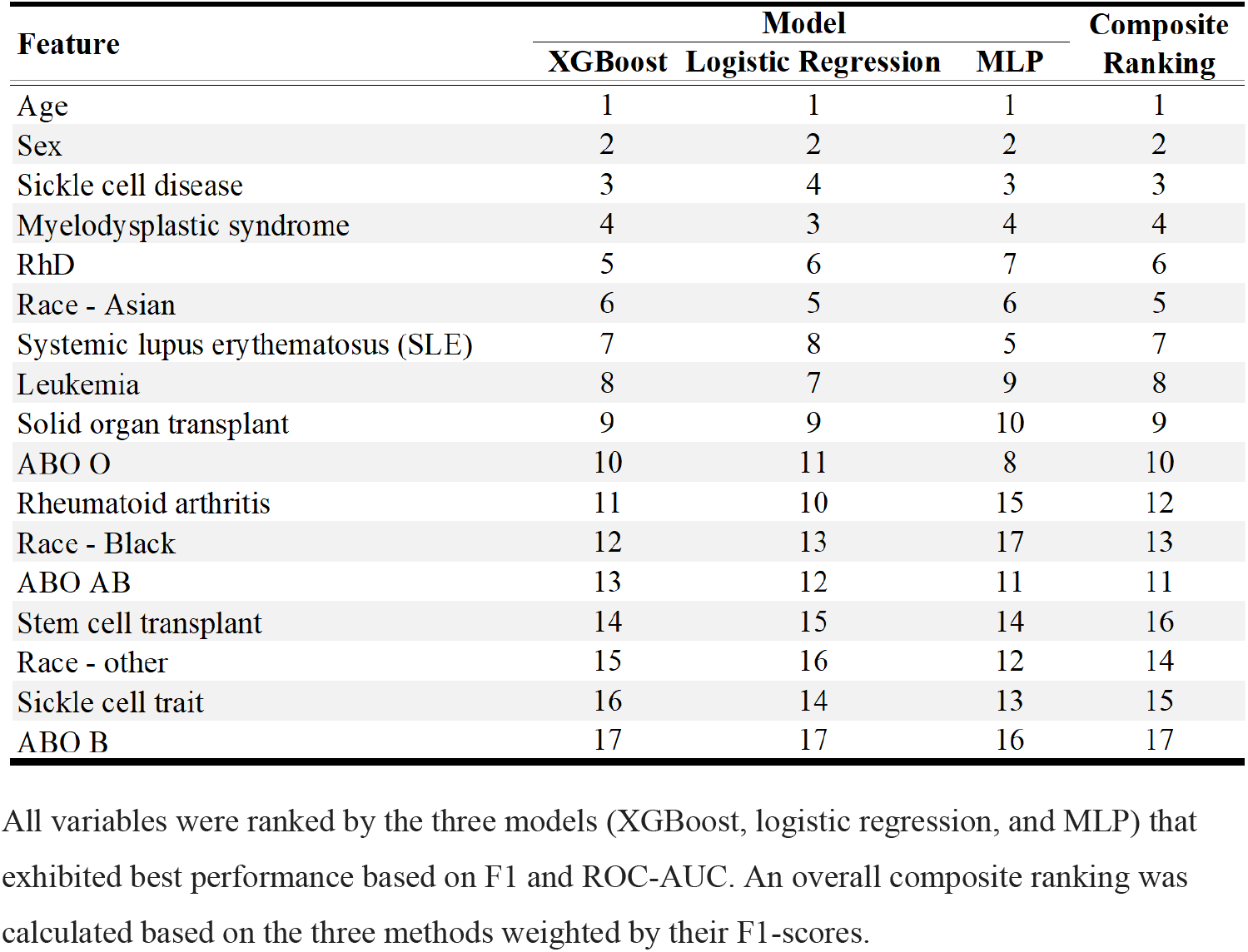
Feature ranking from the top three models.

| Feature | Model |  |  | Composite Ranking |
| --- | --- | --- | --- | --- |
|  | XGBoost | Logistic Regression | MLP |  |
| Age | 1 | 1 | 1 | 1 |
| Sex | 2 | 2 | 2 | 2 |
| Sickle cell disease | 3 | 4 | 3 | 3 |
| Myelodysplastic syndrome | 4 | 3 | 4 | 4 |
| RhD | 5 | 6 | 7 | 6 |
| Race - Asian | 6 | 5 | 6 | 5 |
| Systemic lupus erythematosus (SLE) | 7 | 8 | 5 | 7 |
| Leukemia | 8 | 7 | 9 | 8 |
| Solid organ transplant | 9 | 9 | 10 | 9 |
| ABO O | 10 | 11 | 8 | 10 |
| Rheumatoid arthritis | 11 | 10 | 15 | 12 |
| Race - Black | 12 | 13 | 17 | 13 |
| ABO AB | 13 | 12 | 11 | 11 |
| Stem cell transplant | 14 | 15 | 14 | 16 |
| Race - other | 15 | 16 | 12 | 14 |
| Sickle cell trait | 16 | 14 | 13 | 15 |
| ABO B | 17 | 17 | 16 | 17 |

All variables were ranked by the three models (XGBoost, logistic regression, and MLP) that exhibited best performance based on F1 and ROC-AUC. An overall composite ranking was calculated based on the three methods weighted by their F1-scores.

**Figure 5.**
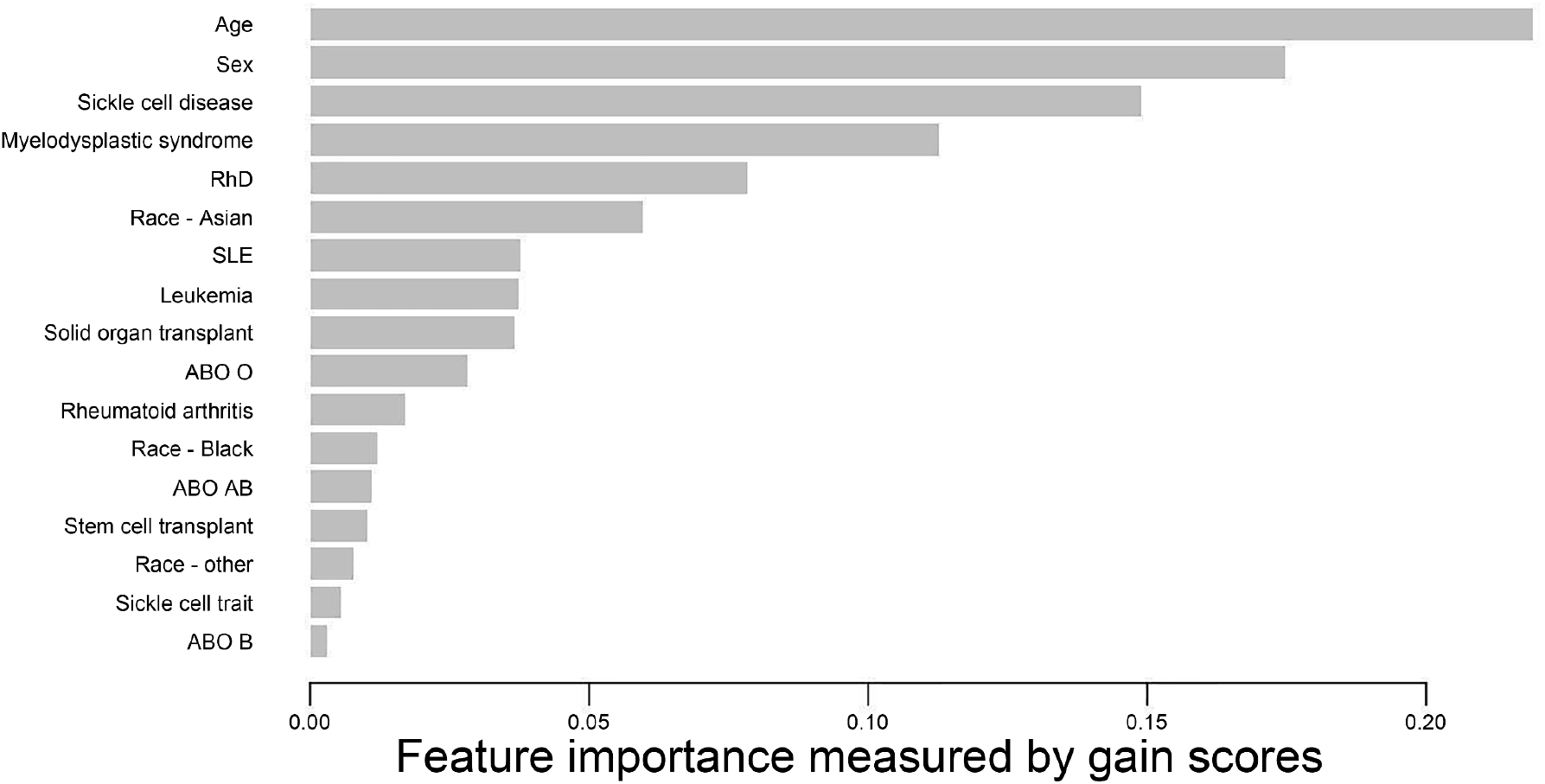
Transfusion Recipient Feature Importance in Alloimmunization. **Ranked importance of predictors in the XGBoost model.** The scores for the predictors are the gain scores from xgb.importance() function. The gain score of a feature measures the fractional contribution of the feature to the mode or the improvement in accuracy brought by the feature, thus quantifying how much the feature contributes to the model’s predictive power, with higher values indicating more important features.

### LLM’s chatbot potential

Although Qwen cannot provide prediction uncertainty directly, it demonstrated considerable potential. One way to obtain estimation uncertainty is to query Qwen models multiple times and calculate the proportions of “yes” prediction to whether a patient will develop alloimmunization after blood transfusion, as shown in **Figure 6A**, where a single risk factor is progressively shifted to a higher risk group. We observe that Qwen7B-qlora model (demonstrating the best F1 score as shown in Figure 3) generates increasing higher proportions of “yes”.

**Figure 6.**
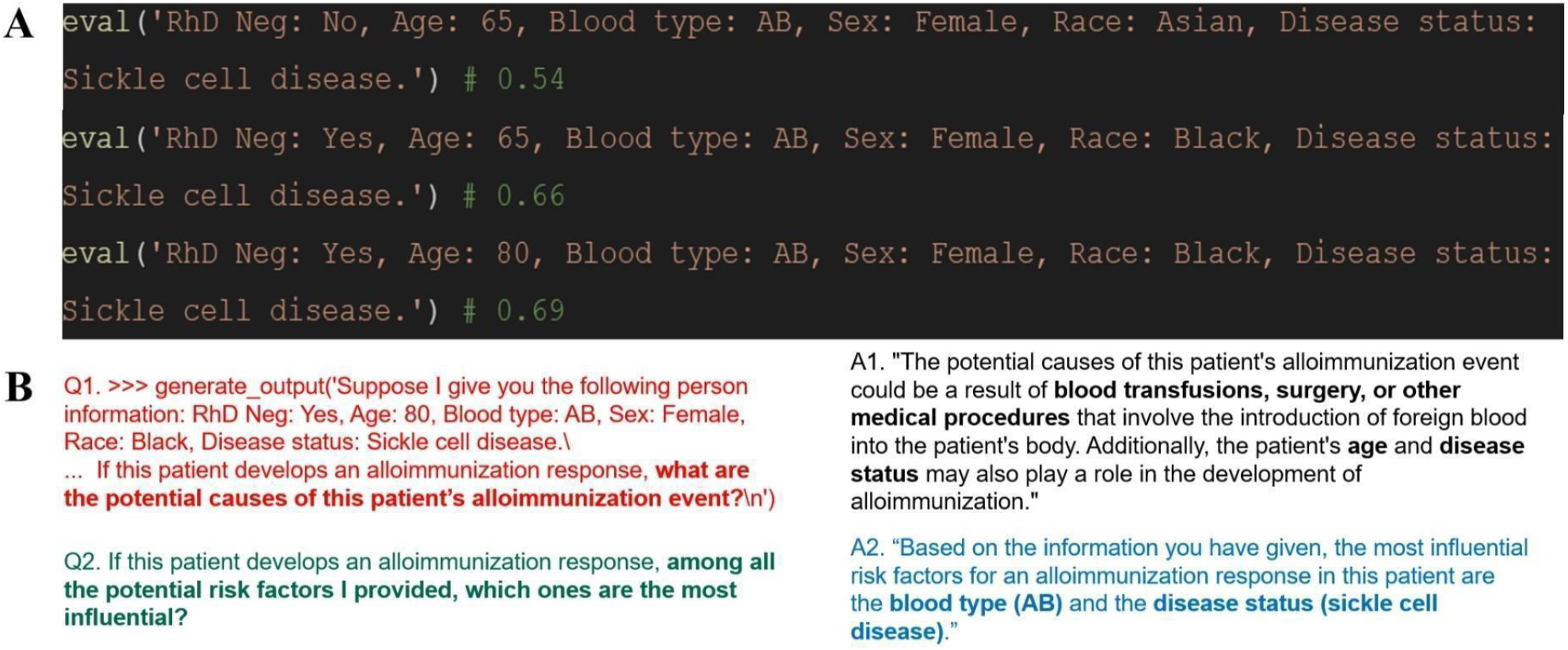
LLM Qwen7B-qlora QA examples. **A**. Risk of RBC alloimmunization for different queries. For each of the three evaluations (note that one feature was changed at a time, to a higher risk group), we entered the prompt 100 times and counted the number of times with a “Yes” answer to whether an alloimmunization event will develop for the patient after blood transfusion. The full prompt is inputs:\nA chat between a curious user and an artificial intelligence assistant. The assistant gives helpful, detailed, and polite answers to the user\’s questions.\nSuppose I give you the following person information: ‘+input_text+’ Does this patient has at least one detectable clinically significant red blood cell alloantibody that could have been induced by transfusion or pregnancy? Use Yes or No to answer.\nAssistant:\n’. **B**. Example QAs to further dive deeper into the causes and most important risk factors.

One advantage of LLM-based methods is their ability to function as a ‘chatbot’ that directly interacts with potential users such as patients considering RBC transfusion. **Figure 6B** shows examples where potential risk factors and the most influential ones can be highlighted based on the input patient information in a QA manner. Such chatbot functionality can help alleviate the burden of insufficient medical personnel while making healthcare more accessible, interactive, and personalized. However, accuracy, fairness, and ethical considerations must be systematically evaluated and ensured (Poulain, Fayyaz, and Beheshti 2024; Haltaufderheide and Ranisch 2024; Xia et al. 2024).

## Discussion

In this study, we applied multiple statistical, machine learning, deep learning and large language model approaches to build models to predict alloimmunization response among patients receiving RBC transfusion. While there is still room for further improvements, this work represents the first comprehensive effort to leverage different categories of methods for alloimmunization risk prediction. In particular, we explored the latest foundation models that have shown great potential in various contexts, including biomedical research (Theodoris et al. 2023; Cui et al. 2024; Fu et al. 2025). Our findings suggest that XGboost and Qwen LLM are the most promising for clinical application, as they demonstrated superior overall prediction performance, high prediction accuracy for individuals predicted with high confidence, and additional value of LLM serving as an interactive chatbot.

Although risk factors have been extensively studied, few models have been developed to predict alloimmunization risk with multiple risk factors considered simultaneously. Limited efforts for prediction purposes focused on specific factors such as the HLA (Geneugelijk, Thus, and Spierings 2014). Our work represents the first effort to explore various classical and state-of-the-art methods, where the predictive models provide either a quantitative probability of alloimmunization risk or a yes/no of an alloimmunization event. Our results provide a baseline for future evaluations by quantifying the model performance in terms of F1 score and AUC-ROC. Additionally, we provided the importance ranking of the features based on multiple prediction models. The top-ranking transfusion recipient features, including age, sex, sickle cell disease, are consistent with the previously reported risk factors with the largest effect sizes. In addition, the ranking may aid in developing a parsimonious prediction model for RBC alloimmunization with optimal performance and in practice, help prioritize data collection.

Several areas exist where future investigations could further improve the current work. First, this work primarily focuses on potential risk factors on the recipient side because our goal was to develop risk prediction models for individuals who are considering blood transfusion, i.e., the potential recipients. However, blood donor characteristics, antigen profiles, and other blood unit modifications (commonly referred to as “component” modifications) have also been reported to influence risk of alloimmunization. Future studies considering these factors are worth exploring. Second, future efforts could consider more complicated models, for example, by incorporating additional risk factors such as the storage duration of transfused RBCs and the number of RBC transfusions (Yu et al. 2025), or by explicitly modeling interactions among the potential predictors. To prevent the initial models from being overly complicated and to retain a larger sample size (since including more risk factors leads to progressive sample size attrition), we decided on the initial set of risk factors in our models. Regarding interaction terms, although not explicitly modeled, XGBoost, MLP and LLMs in theory allow non-linear patterns and may have captured such higher-order relationships. Lastly, and importantly, independent validation in cohorts beyond REDS-III will be highly valuable for assessing the generalizability of the predictive models.

Although further investigations are warranted, our study represents the first comprehensive effort to deploy various statistical, machine learning, deep learning and large language models for the prediction of RBC alloimmunization events. Our study quantifies prediction performance, provided a baseline benchmark for future research, and highlights the model’s potential for individuals predicted with high confidence.

## Data Availability

The data used in this study are available through BioLINCC with Accession Number HLB02071919a.

## Acknowledgements

We thank the REDS-III participants and BioLINCC for granting us data access.

## Supplementary Materials

### Association details

Our association results are consistent with our previous findings (Karafin et al. 2018). Specifically, RhD positive individuals are found to be associated with lower risk of developing RBC alloimmunization (OR = 0.731, P value < 0.0001), and Asians at lower risk (OR = 0.462 and P value < 0.0001), while females (OR = 1.491, P value < 0.0001) and O blood type individuals (OR = 1.145, P value = 3.39×10–4) are at higher risk. We also observed a clear trend suggesting elevated risk for older recipients. For instance, individuals over 40 years old had a 2.6 to 3.7 fold increased risk to be a responder compared with the study referent group (individuals less than 1 year old) with P value ranging from 2.11×10–4 to 4.88×10–5. Individuals with existing disease diagnosis are generally associated with increased risk of RBC alloimmunization. Among them, individuals with sickle cell disease and MDS exhibited the highest odds of being a responder with OR = 6.482 (P value < 0.0001) for sickle cell disease, and OR = 2.193 (P value < 0.0001) for MDS, respectively.

## Supplementary Tables

**Supplementary Table 1.** ICD9/10 codes used in the study.

| Disease | ICD9/10 codes |
| --- | --- |
| leukemia | ICD 9: 204.x*; 205.x; 206.x; 207.x; 208.x<br>ICD 10: C91.x - C95.x |
| malignancy | ICD 9: 140.x - 199.x; ICD 10: C00.x - C80.x |
| myelodysplastic syndrome | ICD 9: 238.7; ICD 10: D46.x |
| rheumatoid arthritis | ICD 9: 714.0, 714.1, 714.2, 714.4, 714.81, 714.89, 714.9<br>ICD 10: M05, M06 |
| sickle cell disease | ICD 9: 282.6, 282.6x. ;<br>ICD 10: D57.0x, D57.1x, D57.2x, D57.8x |
| sickle cell trait | ICD 9: 282.5; CD 10: D57.3 |
| solid organ transplant | ICD 9: V42.x (x=0-7,84,83)**; ICD 10: Z94.x (x=0-7,82, 83) |
| stem cell transplant | ICD 9: V42.83, ICD 10: Z94.84 |
| systemic lupus erythematosus (SLE) | ICD 9: 710.0(sle), 710.1(discoid)<br>ICD 10: M32.0, M32.1, M32.8, M32.9(sle), L93.0(discoid) |

\* 'x' represents all secondary codes unless explicitly stated otherwise.
\*\* Note that, to ensure solid organ transplant, we did not include unspecified transplant, specifically V42.89 and V42.9, which are 'Other specified organ or tissue replaced by transplant' and 'Unspecified organ or tissue replaced by transplant', respectively.

## Supplementary Figures

**Supplementary Figure 1.**
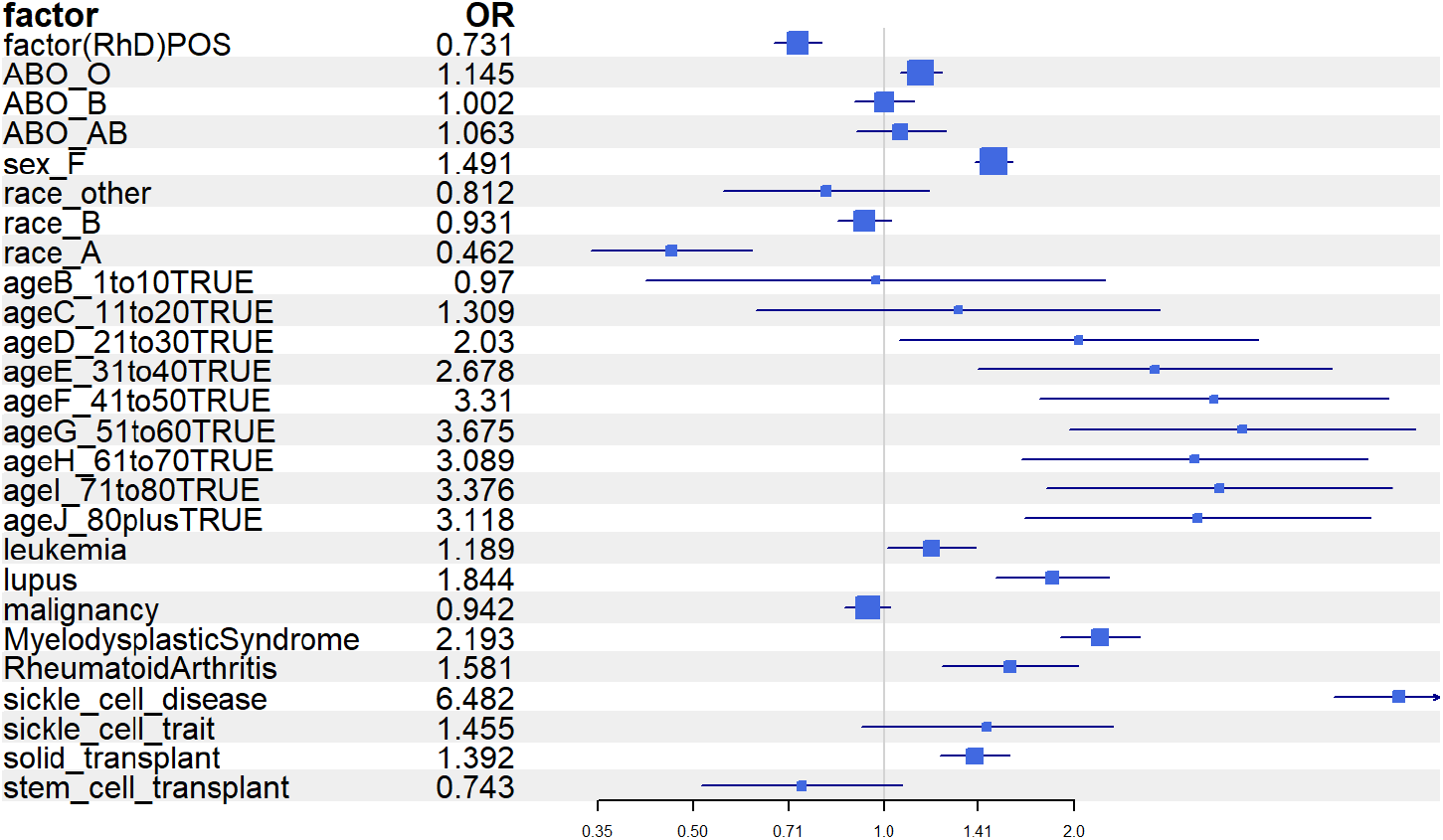
Forest plot for logistic regression results. Size of the blue box indicates the magnitude of OR and the whiskers denote the 95% confidence interval (CI).

**Supplementary Figure 2.**
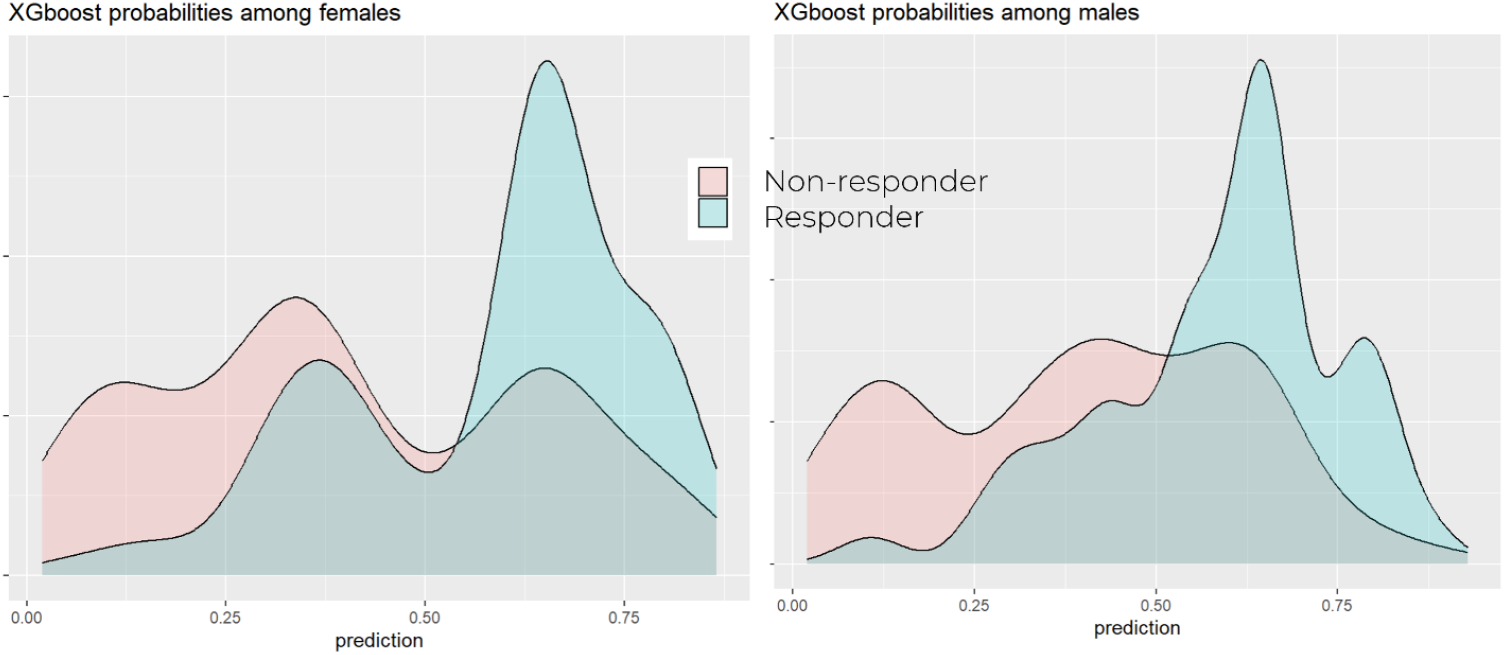
XGBoost model performance for males and females. Density plot for XGBoost predicted probability of being a responder is shown separately for responders (cyan) and non-responders (red), for females (left panel) and males (right panel).

**Supplementary Figure 3.**
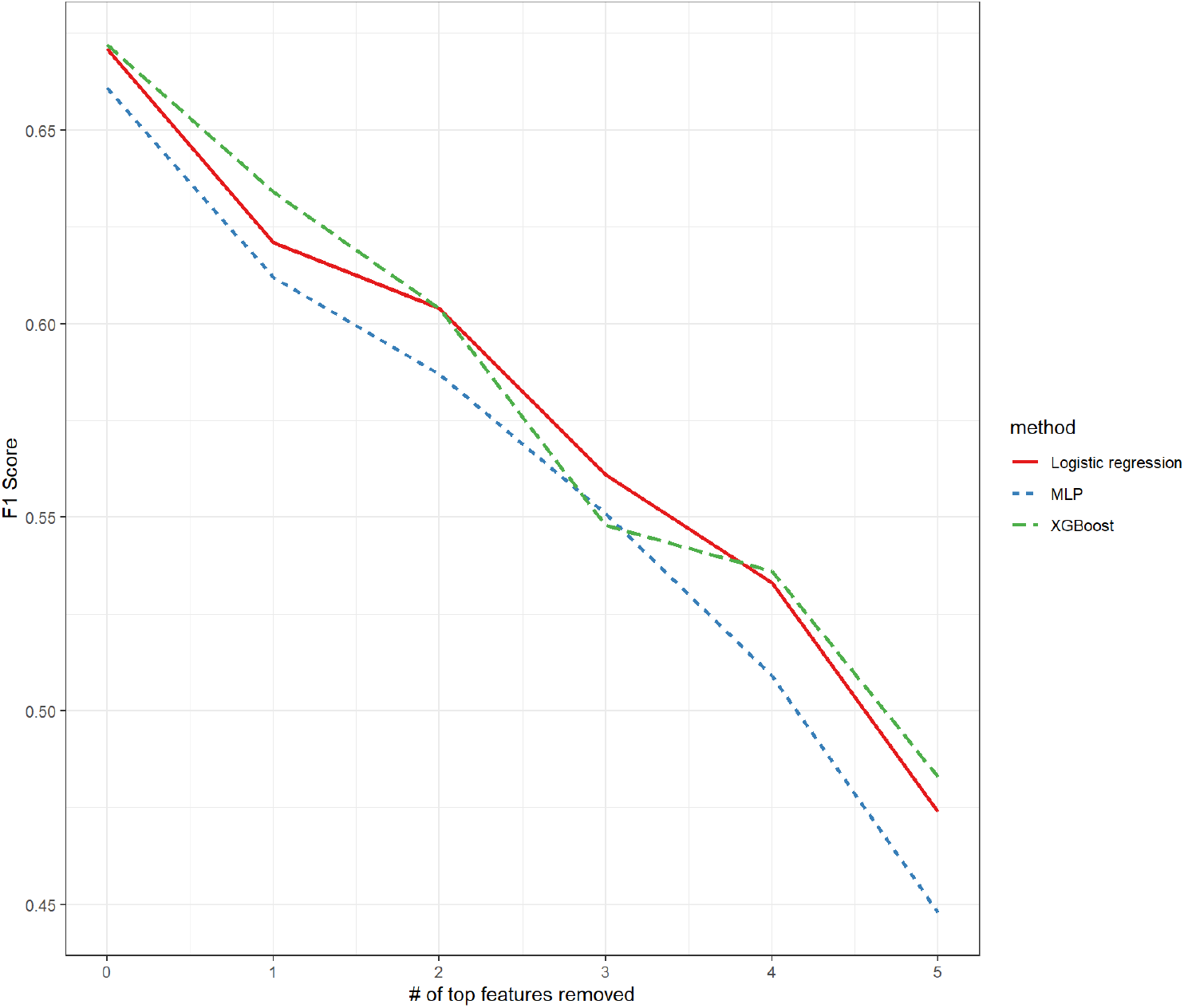
Impact of top features on F1 score. We progressively removed top ranking features for each model and calculated F1 score after feature removal.

**Supplementary Figure 4.**
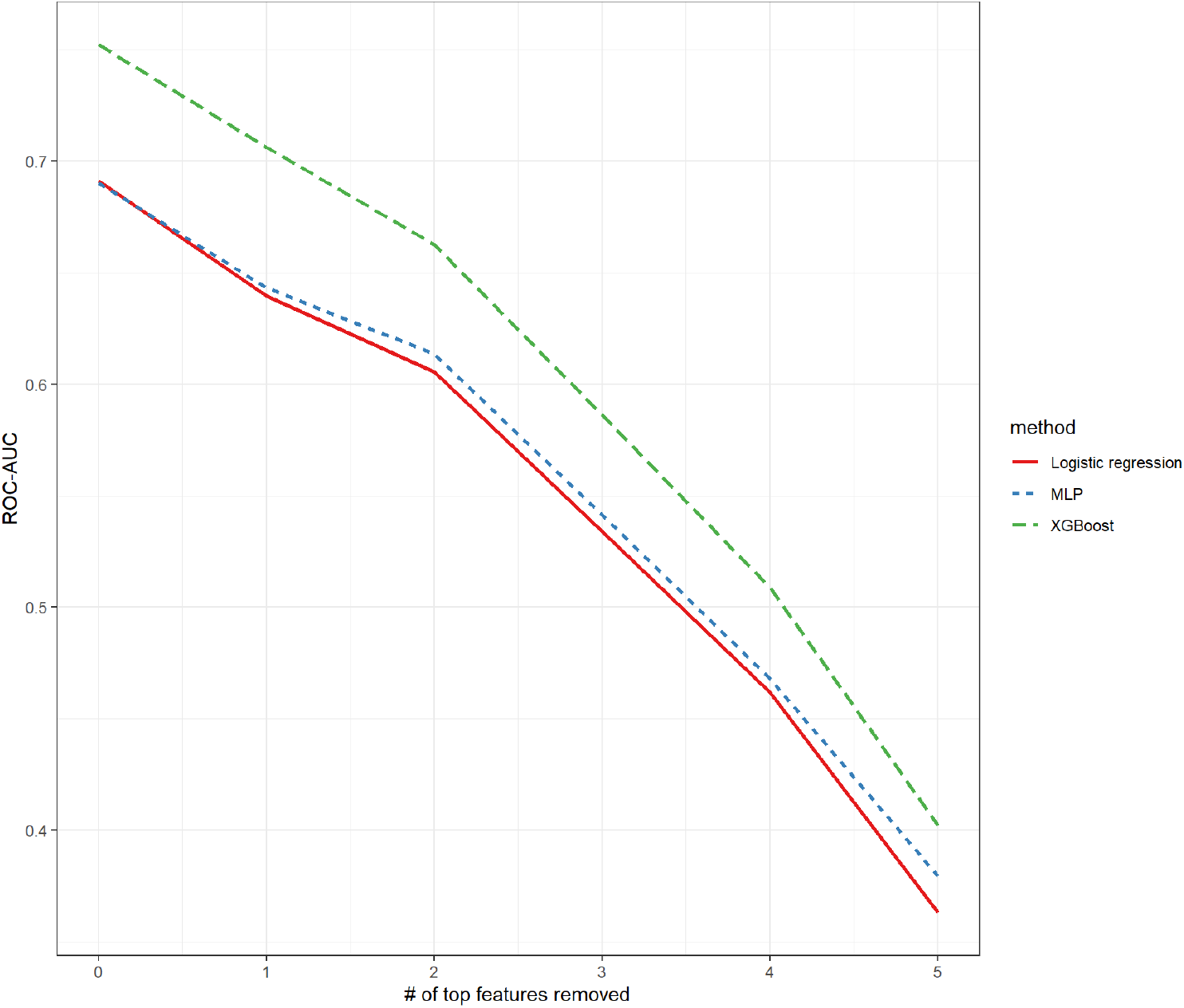
Impact of top features on ROC-AUC. We progressively removed top ranking features for each model and calculated ROC-AUC after feature removal.

